# Factors influencing riverine utilization patterns in two sympatric macaques

**DOI:** 10.1101/2020.06.11.145409

**Authors:** Yosuke Otani, Henry Bernard, Anna Wong, Joseph Tangah, Augustine Tuuga, Goro Hanya, Ikki Matsuda

## Abstract

Many species of terrestrial animals, including primates, live in varied association with the aquatic (e.g., riverine or coastal) environment. However, the benefits that each species receive from the aquatic environment are thought to vary depending on their social and ecological characteristics, and thus, elucidating those benefits to each species is important for understanding the principles of wild animal behaviour. In the present study, to gain a more complete picture of aquatic environment use, including social and ecological factors in primates, factors affecting riverine habitat utilization of two macaque species *(Macaca nemestrina* and *M. fascicularis)* were identified and qualitative comparisons were made with sympatric proboscis monkeys *(Nasalis larvatus)*, which have different social and ecological characteristics. Temporal variation in sighting frequency of macaques at the riverbanks was positively related to the fruit availability of a dominant riparian plant species and negatively related to the river water level which affects the extent of predation pressure. Riverine utilization of macaques was greatly influenced by distribution and abundance of food (especially fruit) resources, possibly in association with predation pressure. Additionally, qualitative ecological comparisons with sympatric proboscis monkeys suggest that the drivers of riverine utilization depend on the feeding niches of the species, and different anti-predator strategies resulting from their differing social structures.

## Introduction

Animals often live in forests characterized as mosaic habitats comprising both terrestrial and aquatic (e.g., riverine or coastal) environments while varying their associations with the aquatic environment. In addition to direct relationships such as obtaining food from the aquatic environment, e.g., bears eating salmon ^1^; racoons eating mussels ^2^, the aquatic environment has an indirect impact on terrestrial animals by bringing about environmental heterogeneity. Differences in terrestrial flora, which are due to varying water abundance and light intensity at the border between aquatic and terrestrial environments, affect foraging behaviour of animals that use plants as a food source. In addition, such boundaries restrict the distribution and movement patterns like a river-barrier ^3^ and thus are constraints on habitat utilization. Aquatic environments are, therefore, undoubtedly important for terrestrial animals across many taxonomic groups and geographic areas.

Primates are a primary example of terrestrial animals that rely on the aquatic environment. Kempf ^4^ presented a comprehensive review of primate aquatic behaviours with the conclusions that the use of aquatic resources and the aquatic environment affect various aspects of primate life, including feeding, traveling, predation avoidance, and thermoregulation, although it has been noted that the represented data are inadequate to draw strong conclusions. Reports of aquatic-related primate ecology have increased in recent years, strongly supporting such proposed broader trends with special reference to primates inhabiting flooded habitats ^5^. One of the notable flooded habitats is riverine forest, providing a relatively constant availability of fruits, combined with greater plant diversity and higher leaf quality when compared with dryland forest, which is due to the frequent supply of nutrient-rich soils in riverine forests that are exposed to seasonal flooding ^6,7^. Further, in contrast to dryland forest, riverine forest potentially has a better light environment and more gap-specialists that have leaves containing higher protein content ^8^—one of the important factors influencing primate abundance ^9^ and dietary choice e.g., ^10,11^. The distinctive food conditions caused by the presence of such large-scale riparian areas in forest habitat are likely to have a significant impact on primate behavioural ecology, and thus riverine habitat is an ideal forest type for research contributing to a fundamental understanding of primate behavioural ecology, including how the aquatic environment affects primate distribution and ranging behaviours.

Furthermore, studying habitats in riparian environments also plays an important role in understanding the landscape of fear ^12,13^. Animals’ responses to predation risk vary over time and space; for example, they can alter their behaviour, such as by changing time allocation patterns depending on the level of fear ^14^. Boundaries between aquatic and terrestrial environments, e.g., riverbanks, can be advantageous or disadvantageous in anti-predator strategies, because they provide animals with a physically heterogeneous environment. In riverine habitats, river-edge trees in exposed places are preferred by primates for roosting to avoid predation, because this allows detection of approaching predators ^15-18^. For example, proboscis monkeys (*Nasalis larvatus*) often choose sleeping sites in river-edge trees in areas with narrow river widths; this provides good escape routes from terrestrial predators such as clouded leopards (*Neofelis diardi*) ^19-21^, which generally show a nocturnal activity pattern ^22^. Conversely, on flooded days when water levels are extremely high, proboscis monkeys remain in inland forest because of reduced predation threat, as terrestrial predators are prevented from foraging on the forest floor ^23^. Therefore, riverine forests have temporal and spatial environmental heterogeneity, and revealing the behavioural changes of animals in response to such heterogeneity can provide insight into behavioural adaptation in response to fear of predators. Consequently, studying primates that live in riverine habitats is ideal for elucidating their resource exploitation patterns relative to anti-predator strategies; these are central components to understanding various primate habitat adaptations.

Long-tailed macaques (*Macaca fascicularis*) and southern pig-tailed macaques (*M. nemestrina*, hereafter pig-tailed macaques, unless otherwise noted) are widely distributed throughout the Sundaic region of Southeastern Asia, often coexisting in a broad variety of habitats ^24^. Long-tailed macaques have often been reported sleeping in river-edge trees ^16,25,26^, and van Schaik, et al. ^16^ indicate that multiple ecological factors (food availability, temperature and predation risk) influence their riverine utilization. Conversely, ecological data for pig-tailed macaques, including the northern species (*Macaca leonina*) and southern pig-tailed species, are less complete ^27-30^, and their aquatic-related behaviours are rarely reported. Rodman ^31^ reported that microhabitat segregation occurs between the two species and that, unlike long-tailed macaques, pig-tailed macaques generally do not use riverine areas. Conversely, Albert, et al. ^18^ noted that the preference of northern pig-tailed macaques to locate their sleeping sites in river-edge trees was a part of their predator avoidance strategy. In addition to predation pressure, as in long-tailed macaques, food availability may affect riverine utilization in pig-tailed macaques. By determining the effects of food availability on riverine forest use in macaque habitat, it is possible to evaluate whether the ranging behaviour of pig-tailed macaques is affected by food availability in the same way as that of long-tailed macaques.

We examined the riverine habitat utilization patterns of sympatric long-tailed and pig-tailed macaques inhabiting a secondary riverine forest on the island of Borneo. At this site, it has been reported that sympatric primates, including the two study macaque species, proboscis monkeys and other species, prefer to utilize river-edge trees for night-time sleeping, although the frequency of riverine usage is different among these sympatric primates ^32^. A previous study on proboscis monkeys at the site suggested that riverine preference is related to high predation pressure, but not river-edge dietary choice ^17,23^. However, in theory, predation risk varies with group size and body weight ^33-35^. The two species of macaques in this study may have different anti-predator strategies from proboscis monkeys, as the macaques live in groups of multiple males and females and have larger group sizes than proboscis monkeys, which live in groups consisting of one male and multiple females ^35,36^ and have significantly different body weight than the two macaque species, i.e., proboscis monkey: male, 25kg and female, 14 kg; long-tailed macaque: male, 6 kg and female, 4 kg; pig-tailed-macaque, male, 14 kg and female, 7 kg ^37-39^. Additionally, the ranging behaviour of primate species that prefer to feed on patchy and clumped food sources, e.g., fruits and flowers, such as northern pig-tailed and long-tailed macaques ^16,25,26^, is more influenced by food distribution and abundance than that of primate species that prefer to feed on ubiquitous food sources (i.e., leaves), such as proboscis monkeys ^40^. Therefore, for macaques, the location of foraging patches may have a stronger effect on sleeping site selection.

In the present study, to gain a more complete picture of the riverine utilization patterns in the two species of sympatric macaques, we sought to 1) evaluate temporal variation in their riverine usage and asses the factors affecting riverine usage, including physical environment, i.e., river width and water level, and 2) describe their diets in river-edge areas, with a comparison of those availability. In addition, the effects of feeding niches and social structure on ranging behaviour are discussed by qualitatively comparing the characteristics of riverine utilization of the two species of macaques with those of proboscis monkeys reported in previous studies ^17,21,23^.

## Methods

### Study area and subjects

We performed the observations over two years from 2012 to 2014 in riverine forests along the Menanggul River (average river width 0–4000 m from the river mouth: 19.9 m), a tributary of the Kinabatangan River, Sabah, Borneo, Malaysia (118°30’E, 5°30’N). The south side of the Menanggul River is covered extensively in natural forest, whereas the north side has been deforested for oil palm plantations, except for a protected zone along the river ^41^. The mean minimum and maximum daily temperatures were approximately 24°C and 30°C, respectively, and the mean annual precipitation at the site was 2,474 mm ^6^. The riverine forest was inhabited by long-tailed and pig-tailed macaques, as well as proboscis monkeys, silver langurs (*Trachypithecus cristatus*), Hose’s langurs (*Presbytis hosei*), maroon langurs (*Presbytis rubicunda*), Bornean gibbons (*Hylobates muelleri*) and orangutans (*Pongo pygmaeus*) ^32^. Long-tailed and pig-tailed macaques under observation were well habituated to observers in boats, as this area is one of the main tourist attractions in the region, with many boats and tourists visiting the Menanggul River since more than 10 years ago.

### Data collection

#### Boat-based surveys

Surveys by boat in the late afternoon are considered the most effective method for studying primates, including the two sympatric macaques in this region, because they often sleep in riverside trees ^32^. We therefore collected data on the distribution pattern of the sympatric macaques in the late afternoon (16:00–19:00) for 434 days from June 2012 to July 2014 via boat-based surveys. We conducted the surveys along the river at a speed of approximately 4–6 km/h. When we detected a group or individual macaque, we switched off the boat engine to avoid disturbing them and paddled closer to record their species and numbers. We divided the river into 50-metre sections from the river mouth to 4,000 m inland, recording the river sections where sightings of macaques were made. When group members were distributed over several sections, the section containing the largest number of individuals in the visual inspection was defined as the detected section.

#### Boat-based behavioural observations

We collected behavioural data from the adults and subadults in both macaque species during the boat-based surveys. During the observation periods, we recorded the activity of all visible primates at the time of detection by scan sampling ^42^ over 60 days from June 2012 to May 2014. We divided the behaviours into seven categories: feeding, grooming, moving, resting, playing, fighting and other. Food items consisted of leaves, fruits, flowers and other items, and food plants were taxonomically identified *in situ*.

#### Vegetation survey

We established a total of 16 transects (200–500 m × 3 m) on both sides of the river at 500 m intervals from the river mouth to 4,000 m. The total length of the 16 transects was 7,150 m, and the total surveyed area was 2.15 ha. We taxonomically identified trees ≥10 cm in diameter at breast height (DBH) and vines ≥5 cm in diameter that were located within the transects ^6^. Because these data did not include herbaceous climbers, we added data on *Cayratia trifolia* (Vitaceae), an important food source for macaque species (see results) in case the climber was entangled in the surveyed trees/vines.

#### *Monthly availability survey for* C. trifolia *fruits*

We carried out a fruit quantity survey of *C. trifolia*, which has fleshy, juicy, dark purple and nearly spherical fruits ca. 1 cm in diameter ^43^. It has been reported that sympatric proboscis monkey at this study site consume *C. trifolia* ^40^. Early in each monthly survey from July 2012 to June 2014, we travelled by boat up to 4000 m from the river mouth and counted all the visible mature and young fruits of *C. trifolia* on both riverbanks. Two observers independently counted the number of fruits in a section, and the average was defined as the availability of *C. trifolia* in the section. We judged the degree of fruit maturation by its colour, with mature fruit being purple to black and young fruit being green.

#### Rainfall, water level and river width

We measured daily rainfall every morning at base camp approximately 1.5 km from the mouth of Menanggul River, using a tipping bucket rain gauge. We also recorded water level and river width to evaluate the effects of river level on behaviours of the study macaques. We installed a water level gauge at the mouth of Menanggul River, and measured the water level at the end of the boat-based survey (17:00–19:00). We measured the river width at the start and end points of each 50 m section with a laser rangefinder, and the average value of the start and end points was used as a representative value of the river width of the section.

#### Data analysis

To extract time-series characteristics of the increase or decrease in the number of *C. trifolia* fruit, we conducted seasonal decomposition of time series by loess (STL) ^44^ which is a filtering procedure for decomposing a seasonal times series into three components: trend, seasonal, and remainder or residual.

We evaluated the effects of the availability of *C. trifolia* fruits, rainfall and river width on the sighting frequency of the two macaque species counted during the boat-based surveys, using a hierarchical Bayesian continuous-time structural equation model (CtSEM). Models were fitted using the R package ctsem ver. 2.5.0 ^45^ with four chains and 4,000 iterations. The CtSEM modelling addresses unequally spaced time intervals in longitudinal data assessment. Since the number of survey days varied between months and the survey days were not evenly spaced, the monthly mean values were not strictly equally spaced time interval data. Through a hierarchical Bayesian framework, CtSEM allows for the estimation of continuous time processes of a sample while accounting for potential subject-level deviations by using population model estimates to inform subject-level model priors ^46^. The possible temporal autocorrelations among our data are the total amount of *C. trifolia* fruits, and the monthly mean number of counted macaques on each day. In addition, since there is a possibility that spatial autocorrelation occurs between adjacent 50 m sections, the 500 m section was adopted for examination of factors for sightings of macaques on riverbanks. As a result of the evaluation of the spatial autocorrelation by Moran’s *I* index, which is the most commonly used coefficient in univariate autocorrelation analyses, such a trend was not detected at each 500 m section for the monthly sighting frequency of the macaques, i.e., mean monthly number of counted pig-tailed (Moran *I* statistic index = −0.324 – −0.115, *p* > 0.3) and long-tailed macaques (Moran *I* statistic standard deviate = −0.299 – −0.085, *p* > 0.2), and availability of *C. trifolia* fruits (Moran *I* statistic standard deviate = −0.224 – −0.101, *p* > 0.2). This indicates that the 500 m section is a unit that can be analysed without considering spatial autocorrelation and is a suitable analytical unit for subsequent analysis. Predictor and independent variables were z-standardized to build a common metric. We performed the calculations using R ver. 3.6.1 ^47^. In the representation of the result of the model, SD refers to posterior standard deviation and PCI refers to posterior credibility intervals. The PCI indicates the probability that the parameter falls between the lower (2.5%) and upper (97.5%) limits.

## Results

### Boat-based survey: sighting frequency of primates and consumed food items

During the study period, there were a total of 3,180 detection events for six species of primates, and 39,907 individuals were observed during boat-based surveys (Table 1). Long-tailed and pig-tailed macaques accounted for 37.0% and 25.7% of the total number of observed individuals, respectively (Table 1). We collected a total of 66 and 277 feeding records for pig-tailed and long-tailed macaques, respectively. Fruits and flowers/buds of *C. trifolia* and *Dillenia excelsa* were by far the most important foods at the riverbanks, which constituted 22.7%, and 50.9% (*C. trifolia*), and 42.4% and 18.8% (*D. excelsa*) of the total feeding records in pig-tailed and long-tailed macaques, respectively (Table 2).

**Table 1.** Summary of detection events and the number of individuals sightings during boat-based surveys (n = 434 days).

|  | Pig-tailed<br>macaque | Long-tailed<br>macaque | Proboscis<br>monkey | Silver<br>langur | Maroon<br>langur | Orangutan |
| --- | --- | --- | --- | --- | --- | --- |
| Total number of discovery<br>event (times) | 442 | 1357 | 1278 | 44 | 1 | 67 |
| Total number of individual<br>sighting (head) | 10257 | 14785 | 14393 | 344 | 6 | 122 |
| Averaged individual<br>sighting (head/day ±SD) | 23.63 ±24.07 | 34.07 ±23.05 | 33.16 ±23.86 | 0.79 ±2.67 | 0.01 ±0.29 | 0.28 ±0.91 |

**Table 2.** Food items and parts consumed by southern pig-tailed and long-tailed macaques with their observed frequency during boat-based surveys.

| Species | Type | Pig-tailed macaque |  | Long-tailed macaque |  |
| --- | --- | --- | --- | --- | --- |
|  |  | Number of observation time | % | Number of observation time | % |
| *Albizia corniculata* | fruit | 1 | 1.52 |  |  |
|  | unkown | 1 | 1.52 |  |  |
| *Antidesma thwaitesianum* | flower |  |  | 10 | 3.61 |
|  | fruit |  |  | 2 | 0.72 |
|  | leaf |  |  | 1 | 0.36 |
| *Baccaurea stipulata* | fruit | 1 | 1.52 |  |  |
| *Cayratia trifolia* | fruit | 15 | 22.73 | 141 | 50.90 |
|  | leaf |  |  | 9 | 3.25 |
| *Dillenia excelsa* | flower | 24 | 36.36 | 45 | 16.25 |
|  | fruit | 4 | 6.06 | 7 | 2.53 |
|  | leaf |  |  | 2 | 0.72 |
|  | unkown |  |  | 1 | 0.36 |
| *Eichhornia crassipes* | stem |  |  | 2 | 0.72 |
| *Ficus* sp. | fruit | 2 | 3.03 | 4 | 1.44 |
| *Gnetum gnemonoides* | fruit |  |  | 1 | 0.36 |
| *Mallotus muticus* | flower | 1 | 1.52 | 3 | 1.08 |
|  | fruit | 3 | 4.55 | 8 | 2.89 |
|  | leaf | 1 | 1.52 | 7 | 2.53 |
|  | unkown |  |  | 1 | 0.36 |
| *Nauclea subdita* | flower |  |  | 1 | 0.36 |
|  | leaf | 1 | 1.52 |  |  |
| *Pternandra galeata* | fruit | 1 | 1.52 | 2 | 0.72 |
|  | leaf |  |  | 2 | 0.72 |
| *Spatholobus cf. macropterus* | leaf |  |  | 1 | 0.36 |
| *Vitex pinnata* | fruit |  |  | 1 | 0.36 |
| *Xylosma sumatrana* | flower |  |  | 5 | 1.81 |
| *Ziziphus bornensis* | fruit | 1 | 1.52 |  |  |
| unkown | flower | 6 | 9.09 | 16 | 5.78 |
|  | fruit | 2 | 3.03 | 4 | 1.44 |
|  | leaf | 2 | 3.03 | 1 | 0.36 |
| Total |  | 66 |  | 277 |  |

### Vegetation characteristics and food availability

We marked 1,645 trees and 497 vines (180 species, 125 genera, 52 families) along our 16 trails (for details, see ^6^). *Cayratia trifolia* was entangled in only four of 1,645 marked trees (0.24%), and was located in well-lit forest gaps caused by fallen trees. Conversely, *C. trifolia* was clearly more abundant along the riverbanks, and was found in 12.5%– 72.5% of all 50-m river sections in each monthly survey. Contrary to the distribution pattern of *C. trifolia, D. excelsa* was more abundant inside the forest: of 98 *D. excelsa* plants in the vegetation transects, 84 (86%) were found in the inland forest (>50 m from the riverbanks). The tendency was the same for *Mallotus muticus* (120 of 149 in the inland forest), which was the third most common plant in feeding records of pig-tailed and long-tailed macaques; *Albizia corniculate* (22 of 28 in the inland forest) and *Ficus* spp. (20 of 23 in the inland forest), which were the fourth and fifth most common in the feeding records of pig-tailed macaques; and *Antidesma thwaitesianum* (27 of 28 in the inland forest) and *Xylosma sumatrana* (53 of 67 in the inland forest), which were the fourth and fifth most common in the feeding records of long-tailed macaques.

The mean monthly number of counted *C. trifolia* fruits on the riverbanks was 23,283.7 (SD ±19,707.3; range 1,844–78,505). The mean monthly numbers of young and mature fruits were 22,795.0 (±19,260.0; 1,806–76,002) and 488.6 (±572.4; 38–2,503), respectively. STL based on the number of counted fruits in each monthly survey clarified that the *C. trifolia* fruit availability seasonally fluctuated (Fig. 1). Fruit availability declined in the middle of 2013 and then increased again, and tended to increase and decrease every 4–6 months.

**Figure 1.**
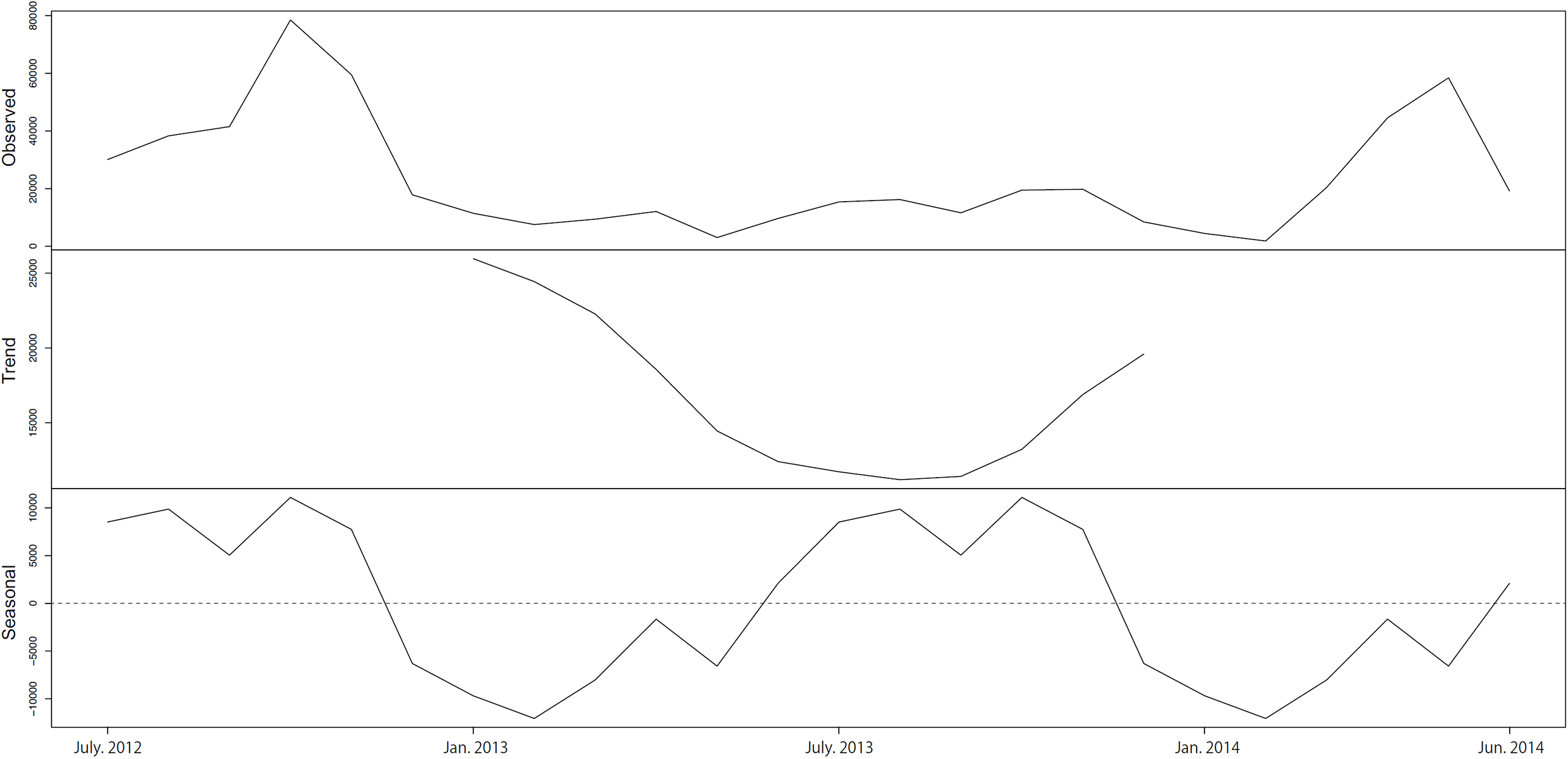
Decomposition plot of abundance of *Cayratia trifolia* fruits based on Seasonal decomposition of time series by loess (STL). Numbers of *C. trifolia* fruits are represented on the y-axis. Trend and seasonality indicate a relatively steady increase or decrease over time, and a pattern that repeats, respectively.

### Factors affecting temporal variation in sighting frequency of macaques

The mean and 95% PCI of T0 mean parameters (Table 3), representing the relationship between the subject’s initial states with their later states throughout the latent process, were 0.408 (−0.089, 0.911), 0.313 (−0.199, 0.783) and 0.270 (−0.230, 0.805) for the sighting frequency of pig-tailed and long-tailed macaques and *C. trifolia* fruits, respectively. This indicated that there was no tendency for each parameter to increase or decrease substantially over time, because for all parameters zero falls within the PCIs. The manifest mean parameters (Table 3) represent the average level of the processes, reflecting the intercepts of the sighting frequency of pig-tailed and long-tailed macaques and availability of *C. trifolia* fruits.

**Table 3.** Means, standard deviations (SD) and posterior credibility intervals (PCI) of the CtSEM model.

| Parameter | Dependent Process |  |  |  |  |  |  |  |  |
| --- | --- | --- | --- | --- | --- | --- | --- | --- | --- |
|  | Mean number of pig-tailed |  |  | Mean number of long-tailed |  |  | Amount of *Cayratia trifolia* fruits |  |  |
|  | mean | SD | PCI<br>[2.5%, 97.5%] | mean | SD | PCI<br>[2.5%, 97.5%] | mean | SD | PCI<br>[2.5%, 97.5%] |
| $T_0$ mean | 0.408 | 0.253 | -0.089, 0.911 | 0.313 | 0.251 | -0.199, 0.783 | 0.270 | 0.262 | -0.230, 0.805 |
| Manifest means | -0.021 | 0.129 | -0.277, 0.233 | -0.030 | 0.078 | -0.192, 0.188 | -0.011 | 0.100 | -0.211, 0.184 |
| **Drift Parameters** |  |  |  |  |  |  |  |  |  |
| Mean number of pig-tailed |  | – |  | 0.984 | 0.367 | 0.356, 1.802 * | 0.557 | 0.446 | -0.264, 1.486 |
| Mean number of long-tailed | 0.753 | 0.281 | 0.272, 1.373 * |  | – |  | 0.101 | 0.215 | -0.309, 0.527 |
| Amount of *Cayratia trifolia* fruits | 1.560 | 0.653 | 0.131, 2.710 * | 0.850 | 0.148 | 0.581, 1.167 * |  | – |  |
| **Effect of time dependent parameters** |  |  |  |  |  |  |  |  |  |
| Rainfall | 0.008 | 0.054 | -0.098, 0.115 | -0.022 | 0.049 | -0.119, 0.075 | 0.020 | 0.053 | -0.084, 0.124 |
| Water level | -0.179 | 0.052 | -0.282, -0.077 * | -0.110 | 0.049 | -0.205, -0.011 * | -0.174 | 0.053 | -0.278, -0.071 * |
| **Effect of time independent parameters** |  |  |  |  |  |  |  |  |  |
| River width | -0.466 | 0.257 | -0.971, 0.045 | -0.155 | 0.259 | -0.638, 0.365 | -0.225 | 0.258 | -0.779, 0.245 |

Asterisks (\*) indicate that zero does not falls within the PCIs, i.e., positive or negative effects are indicated.

The regression coefficients of the monthly mean sighting frequency of macaques and monthly availability of *C. trifolia* fruits within each section denoted the temporal autoregressive effects (Fig. 2); *C. trifolia* fruits had a temporal autocorrelation that lasted approximately three months, but sighting frequency of macaques had no such autocorrelation.

**Figure 2.**
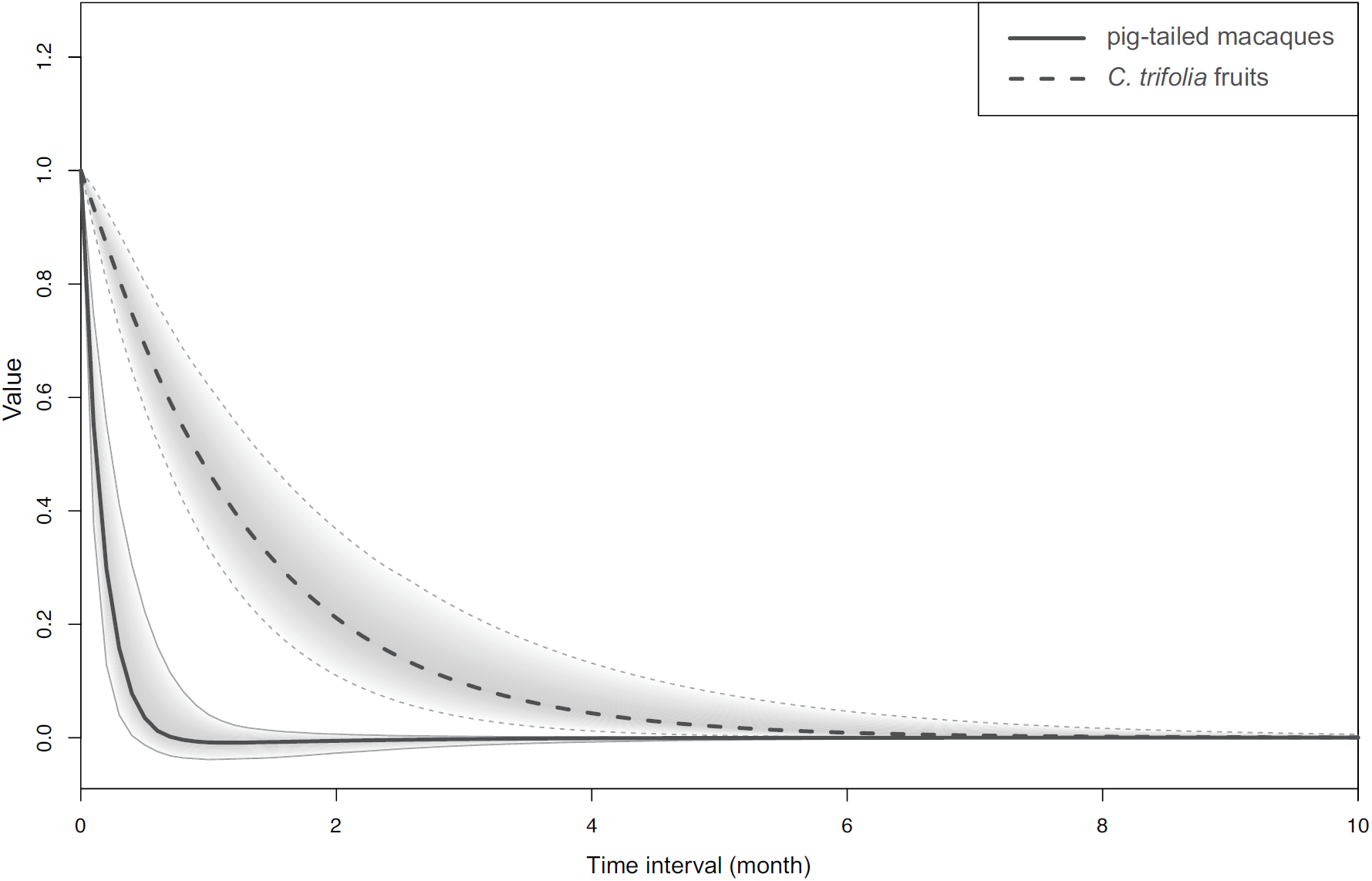
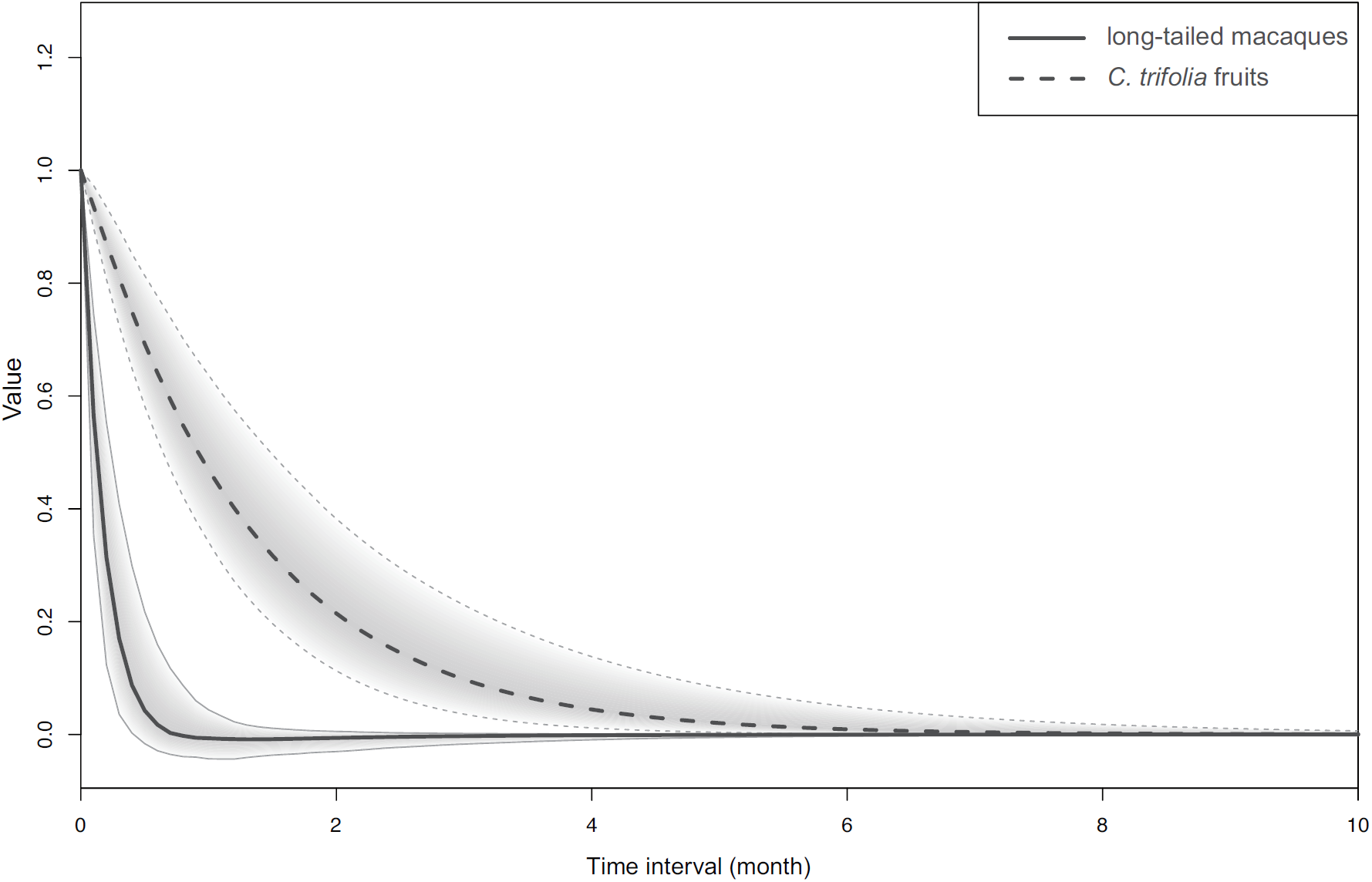
Auto-regressive effects plotted for time intervals of 0 < Δt < 10 months. Parameters represent within-section persistence of the number of *Cayratia trifolia* fruits, and the sighting frequency of pig-tailed and long-tailed macaques over time. Solid lines represent auto-regressive effects for the number of *C. trifolia* fruits over time, and dashed lines represents auto-regressive effects for the sighting frequency of pig-tailed (a) and long-tailed macaques (b).

To assess the effects of the monthly availability of *C. trifolia* fruits on the monthly mean sighting frequency of pig-tailed and long-tailed macaques, we evaluated the drift parameters representing the cross effects (Table 3). The positive values showed that the *C. trifolia* fruits had a positive effect on the sighting frequency of pig-tailed (mean = 1.560, SD = 0.653, PCI = [0.131, 2.710]) and long-tailed (mean = 0.850, SD = 0.148, PCI = [0.581, 1.168]) macaques throughout the study period, indicating that more macaques were sighted in areas where *C. trifolia* fruits were abundant. Conversely, the sighting frequency of macaques did not affect the availability of *C. trifolia* fruits. Additionally, the positive values indicated that the sighting frequency of both macaques had positive effects on each other (pig-tailed to long-tail macaques, mean = 0.984, SD = 0.367, PCI = [0.356, 1.802]; long-tailed to pig-tailed macaques, mean = 0.753, SD = 0.281, PCI = [0.272, 1.373]).

Of the time-dependent/independent variables, we detected neither significantly positive nor negative effects of the monthly rainfall or river width on the mean monthly sighting frequency of both macaque species and the availability of *C. trifolia* fruits, while the water level had a negative effect on those factors (Table 3).

## Discussion

We found that ecological factors influenced the riverine habitat utilization of the two sympatric macaque species in this study. One of the important factors was the availability of *C. trifolia*, which was also the most consumed plant species by the macaques; their temporal variation in sighting frequency at the riverbanks was positively related to the abundance of *C. trifolia* fruits. Conversely, a previous study of proboscis monkeys at this study site reported that food availability is not a fundamental factor for explaining their riverine habitat utilization patterns in the late afternoon ^17^. Differing dietary preference in relation to the digestive physiology between the two macaque species and proboscis monkeys may have created this inconsistency. Hindgut-fermenting primates (e.g., macaques) generally show a stronger preference for fruits than foregut-fermenting primates (e.g., proboscis monkeys), allowing the latter to exploit a diet of leaves in greater quantities ^48,49^. Indeed, *C. trifolia* fruits are not the most preferred food by proboscis monkeys at this study site ^40^.

Predation pressure may also be a factor that affects riverine habitat utilization patterns for the two macaque species in this study, which was also shown for proboscis monkeys ^17^. The landscape of fear is an important driver of prey habitat utilization ^50^. Although it is generally difficult to evaluate predation pressure on primates because of how rare it is to directly observe cases of attempted or successful predation ^34,51^, behavioural responses to predator presence were reported to have more far-reaching consequences for prey ecology than the actual killing of individuals ^12^. Indeed, despite the abundance of food resources when studying these macaques, their use of riverbanks was mainly limited to the late afternoon ^17,32^; one reason for this could be related to their nocturnal anti-predator strategy.

According to previous reports ^20,25,30,52-54^, clouded leopards (*Neofelis diardi*), crocodiles (*Crocodylus porosus* and *Tomistoma schlegeli*), and pythons (*Python* spp.) may be significant potential predators of macaques of any age or sex at this study site. It was previously reported that proboscis monkeys were attacked by clouded leopards when they were in trees ^19^. Therefore, the studied macaques should also be exposed to a threat of predation by clouded leopards, which generally show a strongly nocturnal activity pattern ^22^, when the macaques sleep in trees during the night time; during this time, there is no predation threat from crocodiles. As pythons also tend to move and search for prey during the night, even in trees ^55^, threats of predation on macaques by pythons may be similar to those by clouded leopards and may be predictable.

Riverine habitat utilization in the late afternoon and during sleeping periods at night provides more effective protection against attacks from terrestrial predators such as clouded leopards, because they can only approach the macaques from the landward side. Indeed, several studies reported the use of riverine refugia by long-tailed and northern pig-tailed macaques ^16-18^, possibly to reduce predation risk from such terrestrial predators. Alternatively, the openness of the river banks may pose another problem: vulnerability to predation by raptors. However, raptors that prey on adult diurnal primates are considered to be absent from Southeast Asia ^56^, probably because there are few large raptor species ^57^. According to literature reviews ^58,59^, in the case of immature primates (e.g., infants and juveniles), potential predators in the study area may include raptors such as black eagles (*Ictinaetus malayensis*), crested serpent-eagles (*Spilornis cheela*), and bat hawks (*Macheiramphus alcinus*) ^19^. However, predation upon any primates by these animals was not seen at the study site; thus, their predation pressure on macaques may be less prominent. Therefore, late afternoon and night-time use of the riverbanks by macaques may be a response to fear of nocturnal predators. Further studies that include longer-term direct observations and nocturnal observations would provide direct data on predation (e.g., capture rate, loss rate of group members, and contextual data such as age, sex, and social status of prey) and degree of fear of macaques (e.g., vigilance behaviour inland and on river banks). Such information could provide explicit insight into the nature of the landscape of fear.

Contrary to the anti-predator strategy observed in proboscis monkeys, in which they select sleeping sites in areas with narrow river widths ^19,20^, river width was not a significant factor that predicted the sighting frequency of macaque species at the riverbanks in this study. Although the two species of macaques were rarely observed swimming in the river during this study, river crossing at narrower river sections have been more commonly observed in proboscis monkeys ^21,60^; this may be why river width was not detected as a significant factor for the macaques. Both macaque species live in larger groups with more males than proboscis monkeys ^35,36^, and larger groups are generally more vigilant and are capable of detecting predators from longer distances, which potentially reduces predation risk ^33,61^. Therefore, the benefit of sleeping on the riverbanks for the macaques may simply be the vantage point for detecting approaching terrestrial predators like clouded leopards and pythons, rather than ease of crossing the river. Additionally, to gain a better understanding of our riverine anti-predator hypothesis, further studies should evaluate the differences and similarities of predation vulnerability levels between the two macaque species in terms of their differences in arboreality/terrestriality levels and riverine utilization frequency on the basis of accurate population density estimates in the habitat. Furthermore, many studies emphasized the importance of food resource proximity for sleeping site choice by various primate species e.g., ^18,30,62,63-66^. We do not deny the possibility that selecting sleeping sites on riverbanks may also have a secondary effect of minimizing the macaques’ foraging and traveling costs by sleeping near their feeding areas (i.e., areas abundant in *C. trifolia*).

Echoing our observation of the negative effect of river level on macaque sighting frequency at the riverbanks, Matsuda, et al. ^23^ noted such an effect in proboscis monkeys at this study site; they suggested that this occurred because of reduced predation threats, as terrestrial predators such as clouded leopards are prevented from foraging by deep water covering the forest floor. For the macaques, the negative effect of river level could be caused by decreased attractiveness of the dominant food resources on the riverbanks. Because *C. trifolia* was mostly distributed along lower parts of the riverbanks at this study site, these plants were under water or near the surface of the river when the water level was high. The macaque species in this study may hesitate to forage for *C. trifolia* fruits under such circumstances because of the risk of aquatic predator attacks, despite the reduced terrestrial predator threat due to high-level river water. Indeed, while feeding on *C. trifolia* fruits on low branches (1–3 m above the river), an adult male long-tailed macaque was preyed upon by an estuarine crocodile (*Crocodylus porosus*) ^20^ at our study site. As a result, the increased risks associated with *C. trifolia* foraging because of aquatic predators may have diminished the value of *C. trifolia* as a food resource, and this led to a low sighting frequency of the macaques. Additionally, high water levels may simply make access to the riverbanks more difficult for both macaque species, although this is unlikely: both macaque species were affected by water level, even though long-tailed macaques are more arboreal than pig-tailed macaques ^67^.

This study showed that riverine utilization by pig-tailed and long-tailed macaques was greatly influenced by temporal variation in food resource abundance and predation pressure. In addition, qualitative comparisons of sympatric proboscis monkeys suggested that the drivers of riverine utilization depend on the feeding niches of the species and variations in how they cope with predation pressure due to differences in behavioural patterns and social structure. The sighting frequency of both macaques in the riverine habitat had a positive effect, and their dietary patterns in the riverine habitat were similar; this indicated that their feeding niche separation is ambiguous, especially on the riverbanks, although microhabitat segregation has been reported in these two closely related, coexisting macaque species in East Kalimantan ^26,31^.

It should be noted, however, that we cannot deny the possibility of the microhabitat segregation of these two macaque species in the inland habitat where they were not observed for this study. Co-occurrence of these two macaque species on the riverbanks at our study site may be due to the distinctive distribution of the food resource *C. trifolia*. In fact, the sighting frequency of the macaques had no effect on the subsequent abundance of *C. trifolia* fruits. This indicates that the fruits were super-abundant; therefore, the effect of foraging by macaques on fruit abundance was nearly negligible. The presence of super-abundant food would mitigate feeding competition and allow the two macaque species to co-occur on the riverbanks. Tracking macaque groups into the inland forest during other times of the day would reveal the importance of riverine habitats in the diet of these species and provide further insights into the mechanisms of coexistence. In addition to feeding competition, future studies should evaluate the effects of competition for sleeping trees to elucidate coexistence mechanisms in sympatric macaques, as we observed the two macaque species both sharing a sleeping tree and competing for a sleeping tree.

## Acknowledgments

We thank the Sabah Biodiversity Centre and the Sabah Wildlife Department for granting permission to carry out this research and our research assistants, especially Shah, Asnih, Jasrudy, and the Sabah Forestry Department for support in the field. We appreciate the support of Dr. K. Fukaya in terms of statistical analyses. We thank Clio Reid, PhD, from Edanz Group (https://en-author-services.edanzgroup.com/ac) for editing a draft of this manuscript. Finally, we thank Dr. Julie Teichroeb (handling editor) and two anonymous reviewers for their constructive comments which improved our manuscript. The animals involved in this study were not handled for this study. This study was performed following the Research Guidelines for the Study of Wild Primates of the Primate Research Institute, Kyoto University and in compliance with applicable Malaysian laws.

## Author contributions

YO, GH and IM conceptualized the initial idea; YO and IM performed the data collections; AT, HB, AW and JT arranged the data collection in the wild; YO performed and interpreted the statistical analyses; YO and IM drafted the manuscript; all authors contributed to the final version of the manuscript.

## Data availability statement

Data in support of the findings of this study are available from the corresponding authors by reasonable request.

